# Selective and robust dopamine detection is enabled by aptamer-SWCNT optical sensors in physiological media

**DOI:** 10.64898/2026.02.12.705330

**Authors:** Maria Celina Stefoni, Hanan Yafai, Amelia Ryan, Atara Israel, Ryan M. Williams

## Abstract

Monitoring dopamine in complex biological environments is essential for understanding neurological disorders and disease diagnosis, though it presents a unique chemical challenge. In this work, we rationally designed several single-walled carbon nanotube (SWCNT)-based near-infrared fluorescent sensors for dopamine using ssDNA aptamers as selective molecular recognition elements. The performance of three dopamine-selective aptamer-SWCNT hybrids and sensitive but non-selective (GT)_10_-SWCNT constructs were evaluated and compared for their magnitude of response, sensitivity, and selectivity to dopamine. We performed these studies in buffer, in complex media with noradrenaline and serotonin, and in synthetic cerebrospinal fluid. We evaluated sensor constructs alone, with heat + divalent cation addition, and with four different molecular passivation agents. Ultimately, sensors passivated with bovine serum albumin (BSA) demonstrated strong selectivity for dopamine relative to noradrenaline, serotonin, and ascorbic acid, with a greater magnitude of response compared to (GT)_10_-SWCNT. Concentration-response curves in PBS, in a serotonin and noradrenaline solution, and artificial cerebrospinal fluid (aCSF) revealed dynamic ranges between 30 and 200 nM, and we found that the response occurs within five minutes. Together, these results demonstrate that dopamine aptamer-SWCNT sensors enable more selective and robust optical detection in complex biological environments.

## Introduction

Dopamine is a neurotransmitter involved in motor control, mood changes, reward pathways, and decision making^1^. Disruption of dopamine signaling is associated with neurodegenerative disorders such as Parkinson’s, which involves the progressive loss of dopamine-producing neurons in the brain^1, 2^. It is also linked to Alzheimer’s disease, as studies have shown that behavioral and cognitive alterations can be related with impaired dopamine synthesis^3^. Moreover, elevated dopamine levels in cerebrospinal fluid have been observed in first-episode psychosis patients, showing a positive correlation between illness severity and dopamine concentration^4^.

Monitoring dopamine levels has the potential to improve our understanding of disease progression in the lab and the clinic^5^. This has driven the development of numerous dopamine sensors, including optical^6,7^ and electrochemical^8-10^ detection platforms. For example, a surface-enhanced Raman spectroscopy (SERS) dopamine detection platform based on a silver-coated zinc oxide nanostructured substrate was developed, which exhibited response within the nanomolar range in under 30 minutes^7^. Another example is an electrochemical sensor incorporating a nitrogen-doped graphene microelectrode with a dopamine-specific aptamer, finding a linear detection range between 1 and 100 µM^9^.

Single-walled carbon nanotubes (SWCNTs) are fluorescent nanomaterials formed by a single layer of graphene in a cylindrical structure^11^. The geometry of this lattice is described by the (*n,m*) index, sometimes referred to as chirality, which determines their electronic and optical properties. SWCNTs have been extensively used for *in vitro* and *in vivo* sensing applications, mainly because they fluoresce in the tissue-transparent near infrared (NIR) region and do not photobleach^12^. Due to their hydrophobic nature, SWCNTs must be functionalized to enable dispersion in aqueous solution. Various agents have been employed for this purpose, including nucleic acids, peptides, surfactants, and polymers^12^. Nucleic acids in the form of aptamers not only stabilize nanotubes but they also interact with target analytes, inducing changes in the emission center wavelength and/or intensity^12^. Thus, SWCNT-aptamer sensors have been used to detect various biomarkers including the inflammatory biomarker cytokine IL-6^13^, the stress hormone cortisol^14^, and the neurotransmitter serotonin^15^.

Because of the importance of studying dopamine, and the benefits of using SWCNT as sensor transducers, there has been substantial work using SWCNT-based dopamine sensors^16-24^. These studies primarily employ SWCNT wrapped with guanine-thymine (GT) ssDNA repeats that have no inherent biological selectivity for dopamine. One study used a corona phase molecular recognition (CoPhMoRe) screen to identify polymer- or ssDNA-SWCNT constructs that respond to dopamine. That work selected (GT)_15_, which exhibited a 58-81% increase in fluorescence upon addition of 100 µM of dopamine^17^. That study, however, demonstrated that the response to structurally similar molecules epinephrine and norepinephrine induced the same response as dopamine. Another study used species-enriched (6,5) SWCNT suspended with (GT)_40_ for dopamine detection, exhibiting a similar response to ascorbic acid^19^. Further studies used (GT)_10_-SWCNT which have stronger sensitivity for dopamine^24^. Although these sensors exhibited a strong response to dopamine, the same GT-SWCNT combinations have also been used to detect hydrogen peroxide^25^ and doxorubicin^26, 27^. The lack of GT-SWCNT selectivity for dopamine represents a substantial limitation in dopamine sensing to date given that some of these or others may coexist with dopamine local environments such as cerebrospinal fluid^28-30^. The same limitation exists *in vitro*, as dopamine release events in neural cell cultures are usually accompanied with serotonin and noradrenaline release^28-30^.

In this work, we sought to increase selectivity of optical sensors for dopamine through rational design by incorporating ssDNA aptamers as rational molecular recognition elements. We evaluated the sensitivity and selectivity of several dopamine-specific ssDNA aptamers encapsulating SWCNT constructs. We used several aptamer sequences, tethering strategies, and surface passivation strategies, comparing these rational design methodologies to the field-standard (GT)_10_-SWCNT. Selective sensor construct performance in complex media, including with competitive neurochemicals and cerebrospinal fluid (aCSF) was studied, obtaining a dynamic range between 30 nM and 200 nM. We found that molecular recognition of dopamine improves both the selectivity and sensitivity of optical dopamine sensing in complex environments compared to generic molecular interactions.

## Methods

### Synthesis of ssDNA-SWCNT sensors and controls

High-pressure Carbon Monoxide (HiPCO) SWCNT (NanoIntegris Technologies, Boisbriand, Quebec) were suspended in solution separately with ssDNA oligonucleotides, including three ssDNA aptamers, (GT)_10_, and a palindromic linker^31^ (**Table 1**) (Integrated DNA Technologies, USA) in a 1:2 mass ratio in 1x phosphate-buffered saline (PBS) as previously described^32^.

**Table 1.**
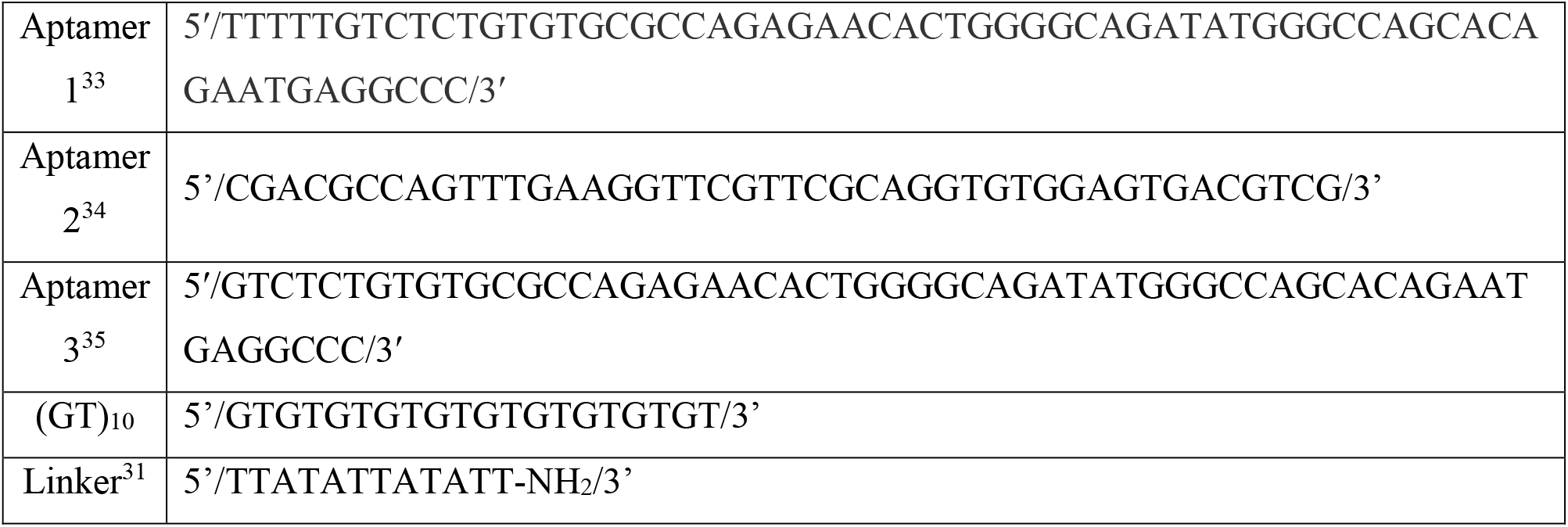
ssDNA sequences used in this study.

Separately, each ssDNA-SWCNT sample was sonicated at 40% amplitude for 60 minutes in an ice bath with a 120 W ultrasonicator and 1/8” probe microtip (Fisher Scientific). The suspension was then ultracentrifuged at 58000 X g for 1 hour (Beckman Coulter) to remove impurities and aggregates. The top 75% of the supernatant was collected. Prior to use, the SWCNT suspensions were filtered through a 100 kDa centrifugal filter (Sigma-Aldrich) to remove excess unbound oligonucleotides and resuspended in 100-200 µl of 1x PBS.

In addition, we synthesized a tethered sequence, in which Aptamer 2 was conjugated to a ssDNA sequence that wrapped the nanotube. This sensor design would decouple the ssDNA suspension from the aptamer binding functions in theory. The palindromic Linker sequence was prepared as above, then 1 µL of Aptamer 2-NHS at a concentration of 10 mg/mL was added to 3 mL of 0.55 mg/L of SWCNT-ssDNA. This was incubated for two hours and then used directly for sensor measurements.

### Characterization of sensor concentration

SWCNT suspended by oligonucleotides were characterized with a V-730 UV-Visible absorption spectrophotometer measured over 300-1100 nm (Jasco Inc.) using the molar extinction coefficient Abs_630_ = 0.036 L mg^−1^ cm^−1^ to determine the concentration of each as previously described^13^.

### High-throughput sensor sensitivity and selectivity

Near-infrared fluorescence spectra of each SWCNT construct and control were acquired via high-throughput NIR spectroscopy (ClaIR, Photon etc., Montreal, Quebec) with laser source excitation wavelengths 655 nm and 730 nm. Near-infrared spectral acquisitions were performed in a Corning half-area UV 96-well plate (Fisher Scientific) with fluorescence spectra acquired between 900 and 1700 nm. Excitation laser power was set to 1750 mW with an exposure time of 200 ms.

All SWCNT fluorescence screening experiments used a nanotube concentration of 0.5 mg/L in 1x PBS and a total volume of 200 µL. A NIR fluorescence baseline was acquired before the addition of the analyte and spectra were obtained at 5 minutes and then every 15 minutes for 3 hours after analyte addition.

To assess sensor selectivity, dopamine, noradrenaline (L-NorAdrenaline 98%, Fisher Scientific), serotonin (Serotonin Hydrochloride 98%, Fisher Scientific) and ascorbic acid (L-ascorbic acid, Fisher Scientific) were added to sensor and control constructs to a final analyte concentration of 100 nM. To assess sensitivity for dopamine and sensor dynamic range, we added 1, 10, 30, 70, 100, 200 and 1000 nM of dopamine in 1X PBS.

We then sought to evaluate selectivity for dopamine in complex environments. First, we added dopamine concentrations of 30, 70, 100, and 200 nM to the sensors in the presence of 50 nM noradrenaline and 50 nM serotonin in 1X PBS. Then, we focused on simulating the environment in which dopamine would ordinarily be found in vivo. We therefore added the same concentrations of dopamine to the sensors in a solution that was complete % artificial cerebrospinal fluid (aCSF, Fisher Scientific, 98% final concentration).

### SWCNT sensor passivation

To prevent nonspecific binding, we investigated several passivation agents in conjunction with the nanosensor constructs. For passivation experiments with sodium cholate (SOC, Fisher Scientific), a final concentration of 7 mM SOC (which is half of the critical micelle concentration, CMC, in PBS^36^) or 3.5 mM (0.25 times the critical micelle concentration) was used with 0.55 mg/L SWCNTs. The mixture was incubated at 4 °C for 12 hours and then incubated with analytes prior to fluorescence spectroscopy. We also performed passivation experiments with sodium dodecyl sulfate (SDS, Fisher Scientific). A final concentration of 0.5 mM SDS (which is half of CMC in PBS^37^) was used with 0.55 mg/L SWCNTs. The mixture was incubated at 4 °C for 12 hours and then incubated with analytes prior to fluorescence spectroscopy.Passivation experiments with an equimolar mix of deoxyribonucleotide triphosphates (dNTPs, Fisher Scientific) were also performed. We added dNTPs at a mass ratio of 50:1 and 100:1 relative to SWCNTs^38^. The solution was incubated at 4 °C for 30 minutes and then incubated with analytes prior to fluorescence spectroscopy.Finally, we performed sensor passivation experiments with bovine serum albumin (BSA, Fisher Scientific), BSA was added at a mass ratio of 45:1 relative to SWCNTs^38^. The solution was incubated at 4 °C for 30 minutes and then incubated with analytes prior to fluorescence spectroscopy.

### Spectral data processing and analysis

All experiments were performed in triplicate. Individual SWCNT (*n,m*) emission peaks were identified according to published studies^13, 32^. Each peak was fit using a pseudo-Voigt model with a custom MATLAB code (available upon request), with data used for analyses when model fit R^2^ was greater than 0.95. Triplicate averages and mean standard deviations were obtained and reported.

Intensity changes were calculated as ((*I* – *I*_0_)/*I*_0_)*100%, which is the difference between the final fluorescence intensity after 3 hours, *I*, and the intensity before the addition of analyte, *I*_0_, (*I* – *I*_0_), normalized by the starting intensity *I*_0_. The results were then normalized by subtracting the sample response to SWCNT intensity in PBS in absence of any analyte. Center wavelength shifts were calculated as the difference in the center wavelength after 3 hours and center wavelength before the addition of analyte (CW - CW_0_). One-way ANOVA plus post hoc Dunn-Sidak tests were performed in OriginLab to determine whether concentration curve values were significantly different from controls.

To describe the binding of dopamine to ssDNA-SWCNT constructs, a Langmuir fit was used. Langmuir fits were used as this binding system is one in which a construct interacts with one analyte and reaches saturation^13^, using the equation:

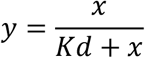

where y is the normalized intensity change, x is the concentration of analyte, and K_d_ is the dissociation constant. Fits were performed with Python code (available upon request) which accounted for variance in mean.

## Results and Discussion

To improve the sensitivity and selectivity of dopamine optical sensors, we investigated four molecularly-specific ssDNA aptamer-SWCNT rational sensor designs, as well as supportive methods to improve selectivity. We compared these to a sensor construct, which was previously published, that has no inherent biological selectivity for dopamine^24^. Of the four rational sensor designs, three consisted of a published aptamer ssDNA sequence directly wrapping SWCNT. The other was an aminated ssDNA sequence (TTA(TAT)_2_ATT)-NH_2_ directly wrapping SWCNT conjugated to an NHS-modified aptamer. Absorbance spectra confirmed effective DNA wrapping for all five ssDNA-SWCNT preparation (**Figure 1A**), and their fluorescence spectra (**Figure 1B**) exhibited intense signals, suitable for sensing applications.

**Figure 1:**
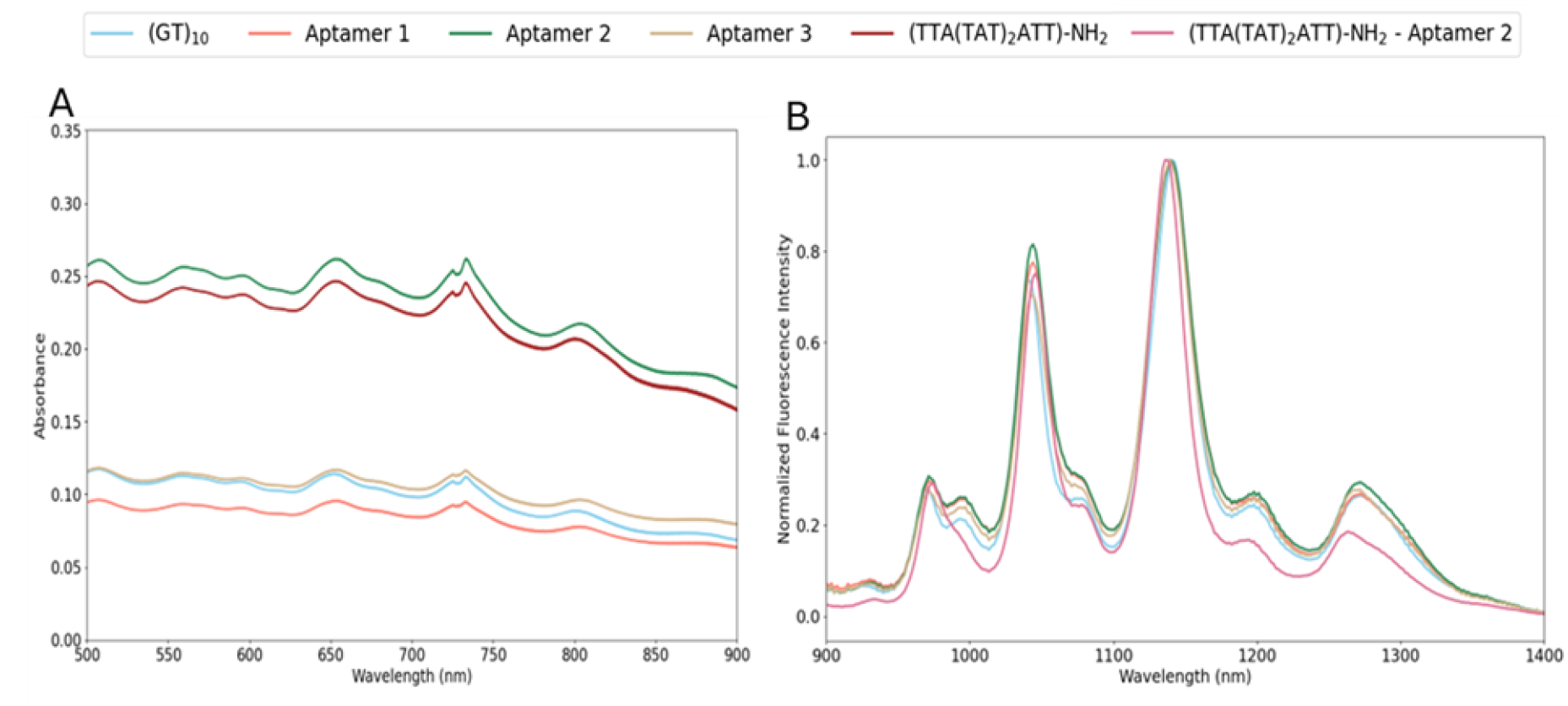
Optical characterization of ssDNA-SWCNT constructs. (A) Absorbance spectra of SWCNT wrapped with (GT)_10_, dopamine Aptamers 1, 2 and 3, and (TTA(TAT)_2_ATT)-NH_2_. (B) Normalized fluorescence intensity obtained using 655 nm excitation laser of SWCNT wrapped with (GT)_10_, dopamine Aptamers 1, 2 and 3, and (TTA(TAT)_2_ATT)-Aptamer 2 conjugate.

### Dopamine aptamer ssDNA-SWCNT sensors selectively detect dopamine

We initially sought to evaluate the selectivity of each molecular recognition element-suspended SWCNT construct. We did so by testing their response, along with (GT)_10_-SWNCT, to dopamine as well as noradrenaline, serotonin, and ascorbic acid. These interfering, or confounding, molecules were chosen due to their structural similarity to dopamine and their coexistence in cerebrospinal fluid and neural cell cultures^28-30^, making selective dopamine detection critical for applications in these complex matrices. We evaluated the addition of 100 nM of each analyte as previous work has demonstrated this to be within the physiologically relevant range^39, 40^.

Both center wavelength shifts (**Supplementary Figure S1A**) and intensity changes (**Supplementary Figure S1B**) of the (7,5) peak were analyzed over a three-hour period. Intensity responses exhibited a relatively more robust response to dopamine and were thus primarily analyzed for these studies. First, we evaluated the (GT)_10_-SWCNT hybrid, finding that only the response to noradrenaline was significantly different than the response to dopamine, whereas very similar responses were found with both serotonin and ascorbic acid (**Figure 2A**). Next, the three reported dopamine-specific aptamer sequences were tested to assess whether the use of a biorecognition specific element could improve selectivity. Indeed, ssDNA aptamer-SWCNT sensors exhibited a selective, and statistically significant, response to dopamine compared to the other three analytes (**Figure 2B)**. Interestingly, each aptamer-based sensor exhibited a larger response to dopamine, with a normalized intensity change of 38%, 29% and 19% for dopamine Aptamers 1, 2 and 3, respectively, compared to just 13% for (GT)_10_. For each aptamer-SWCNT sensor, the response to dopamine was significantly greater than the response to any of the three non-target molecules. In each case, noradrenaline induced little to no response, or a decrease in intensity for Aptamer 3-SWCNT. Ascorbic Acid induced no response for Aptamer 3-SWCNT, and approximately half the intensity increase induced by dopamine for Aptamers 1 and 2. Serotonin induced the greatest intensity increase for all three aptamer constructs.

**Figure 2:**
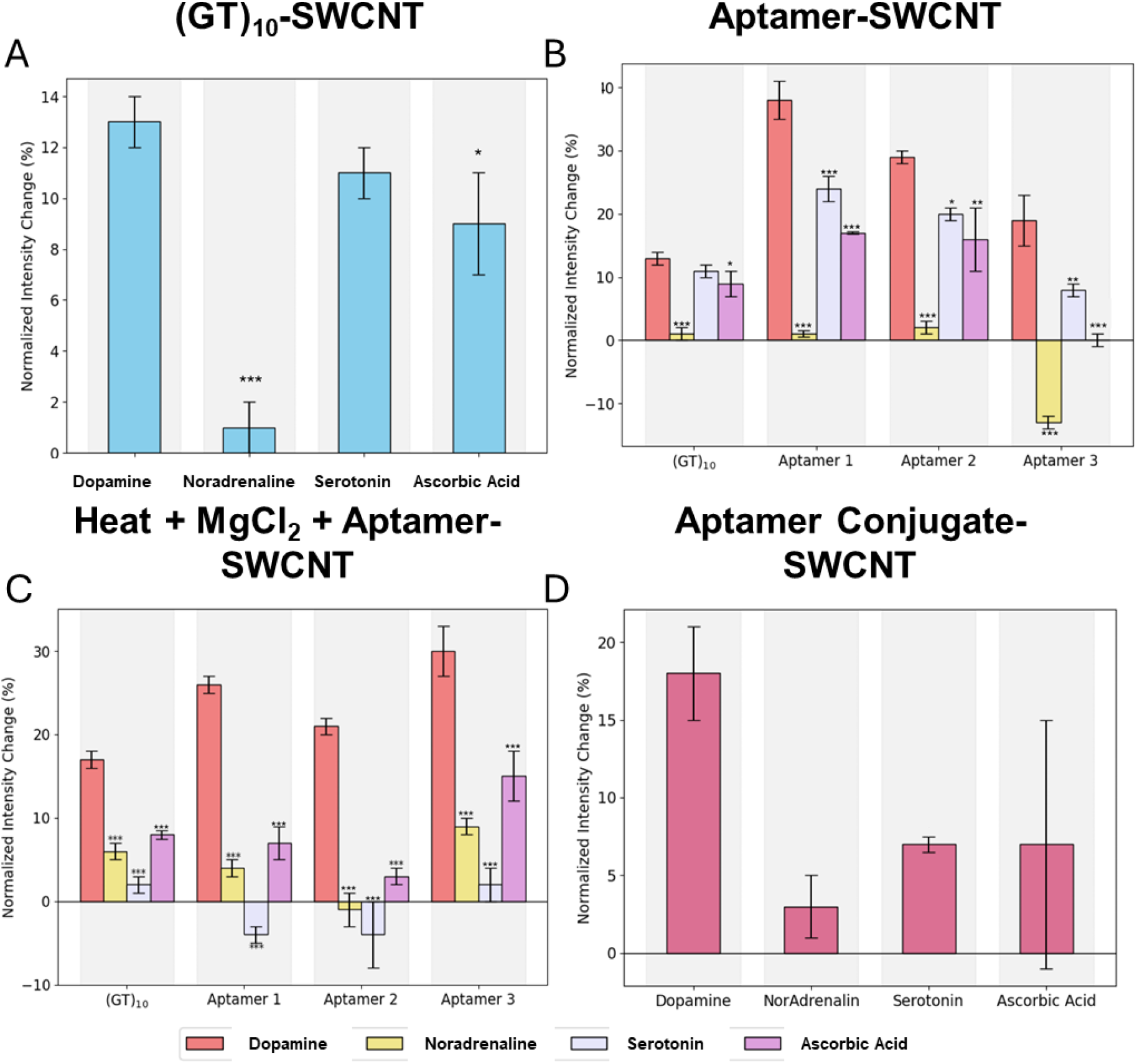
Aptamer-SWCNT sensor constructs detect dopamine more robustly and more selectively than (GT)_10_-SWCNT. Change in fluorescence intensity of SWCNT wrapped with (A) (GT)_10_, (B) three dopamine aptamers, (C) three dopamine aptamer-SWCNT heated to 90°C for five minutes followed by addition of added MgCl_2_, and (D) (TTA(TAT)_2_ATT)-Aptamer 2 conjugate-SWCNT. Dopamine, noradrenaline, serotonin, and ascorbic were added at 100 nM for each experiment. Significant differences in means were determined for a given DNA - SWCNT hybrid by comparing responses to each interferent with those to dopamine, * p<0.05, ** p<0.01, *** p<0.001.

Despite the 1.5 to 3-fold larger magnitude of response for each aptamer sensor, and a significantly larger response to dopamine compared to each interferent, responses to other analytes indeed remained substantial. We chose to use these aptamers in our sensor design as each demonstrated prior selectivity against the interferents used in this study: Aptamer 1 exhibited selectivity against ascorbic acid^33^, Aptamer 2 showed selectivity against serotonin and noradrenaline^34^, and Aptamer 3 was selective against ascorbic acid^35^. Hence, these results suggest that there may be some negative interference due to the dual functionality in this design, including dopamine binding and SWCNT suspension. To promote each aptamer folding into its native conformation, sensor constructs were resuspended in PBS with the divalent cation MgCl_2_, known to be important for proper aptamer folding^41^, after centrifugation and then heated for 5 minutes at 90°C. Upon treatment, the intensity changes with dopamine (**Figure 2C)** were 12% and 8% smaller for Aptamers 1 and 2, respectively, and 4% and 11% larger for (GT)_10_ and Aptamer 3, respectively. Notably, the response to the interferents was smaller than to dopamine with all SWCNT constructs, including (GT)_10_. For instance, for the conformation-induced Aptamer 1-SWCNT construct, the response to dopamine was an increase in fluorescence of 26%, while for noradrenaline, serotonin and ascorbic acid it was of 4%, −4% and 7%, respectively. Without treatment, the response of this construct to these interferents was 1%, 24% and 17%, respectively, compared to 38% response to dopamine, thus suggesting an improvement in selectivity against serotonin and ascorbic acid. Indeed, in the appropriate conformation, Aptamer 2 demonstrated strong selectivity for dopamine compared to all three analytes, which is promising given its prior demonstration to be selective against noradrenaline and serotonin^34^.

To further study the impact of ssDNA aptamer conformation and its interactions with SWCNT on dopamine recognition, we decoupled the ssDNA suspension from the dopamine binding functions of the aptamer. This design could prevent the recognition sequence from interacting tightly with SWCNT, allowing it to remain free for analyte recognition. SWCNTs were first dispersed with a ssDNA sequence ((TTA(TAT)_2_ATT)-NH_2_) that has shown strong affinity for SWCNT^31^ but has no inherent biological affinity for dopamine. Aptamer 2 was then conjugated to this sequence, as it was the only one among the three used in this work reported to exhibit selectivity against both serotonin and noradrenaline^34^. This strategy resulted in an 18% increase in fluorescence intensity in response to dopamine, which was significantly greater than minimal responses to serotonin and noradrenaline, as expected (**Figure 2D**). The response to ascorbic acid was somewhat variable. However, this tethered conjugate approach resulted in an increase in intensity which was 20% less than Aptamer 2 alone (**Figure 2B**). This may be attributed to the distance of the SWCNT surface to the aptamer-dopamine binding events, which could reduce the extent of the fluorescence modulation.

### Passivation of the SWCNT surface improves sensitivity and selectivity of aptamer-SWCNT

While the use of aptamers indeed improved dopamine selectivity as well as the magnitude of response, they remained responsive to closely-related analytes. This could be either due to interactions of ssDNA with the interferents or due to non-specific adsorption of the molecules onto the SWCNT surface. To avoid non-specific adsorption, we used molecular passivation agents to assess their ability to block the nanotube surface^38, 42-45^.

We studied four passivation agents to investigate their ability to block nonspecific adsorption of analytes to the SWCNT surface: the bile salt anionic surfactant sodium cholate (SOC), the anionic surfactant sodium dodecyl sulfate (SDS), free DNA bases as deoxynucleotidyl triphosphates (dNTPs), and the globular protein bovine serum albumin (BSA). SOC passivation, using a final concentration that is half of the critical micellar concentration led to a decrease in intensity change compared to controls (**Supplementary Figure S2A**), opposite to the trend previously observed. More importantly, it impaired sensor selectivity. SOC as a passivation agent was also tested at a lower concentration of 0.25 CMC, which again yielded generally poor selectivity for dopamine (**Supplementary Figure S2B**). We chose concentrations below the SOC CMC, though it is possible that SOC still inhibited direct interaction of the aptamer and dopamine, or it impaired aptamer stability. Next, we assessed the passivation potential of an excess of free DNA bases (dNTPs) because of their known π-stacking with graphitic surfaces. At a concentration of 50X dNTPs, response to noradrenaline was stronger than to dopamine **(Supplementary Figure S3)**. However, at concentration of 100X, dopamine response was significantly greater than others, though the overall magnitude was diminished.

We also assessed SDS as a passivation agent, finding that it actually induced a large intensity decrease for all three dopamine aptamer constructs **(Figure 3A)**. It is surprising that SDS passivation induced an increase in intensity in response to all three interferents, and more so that it did not affect the intensity increase in response to (GT)_10_. These results are worth further exploration, though we sought to engineer a ‘turn-on’ sensor rather than a ‘turn-off’ sensor, thus we continued further exploration.

**Figure 3:**
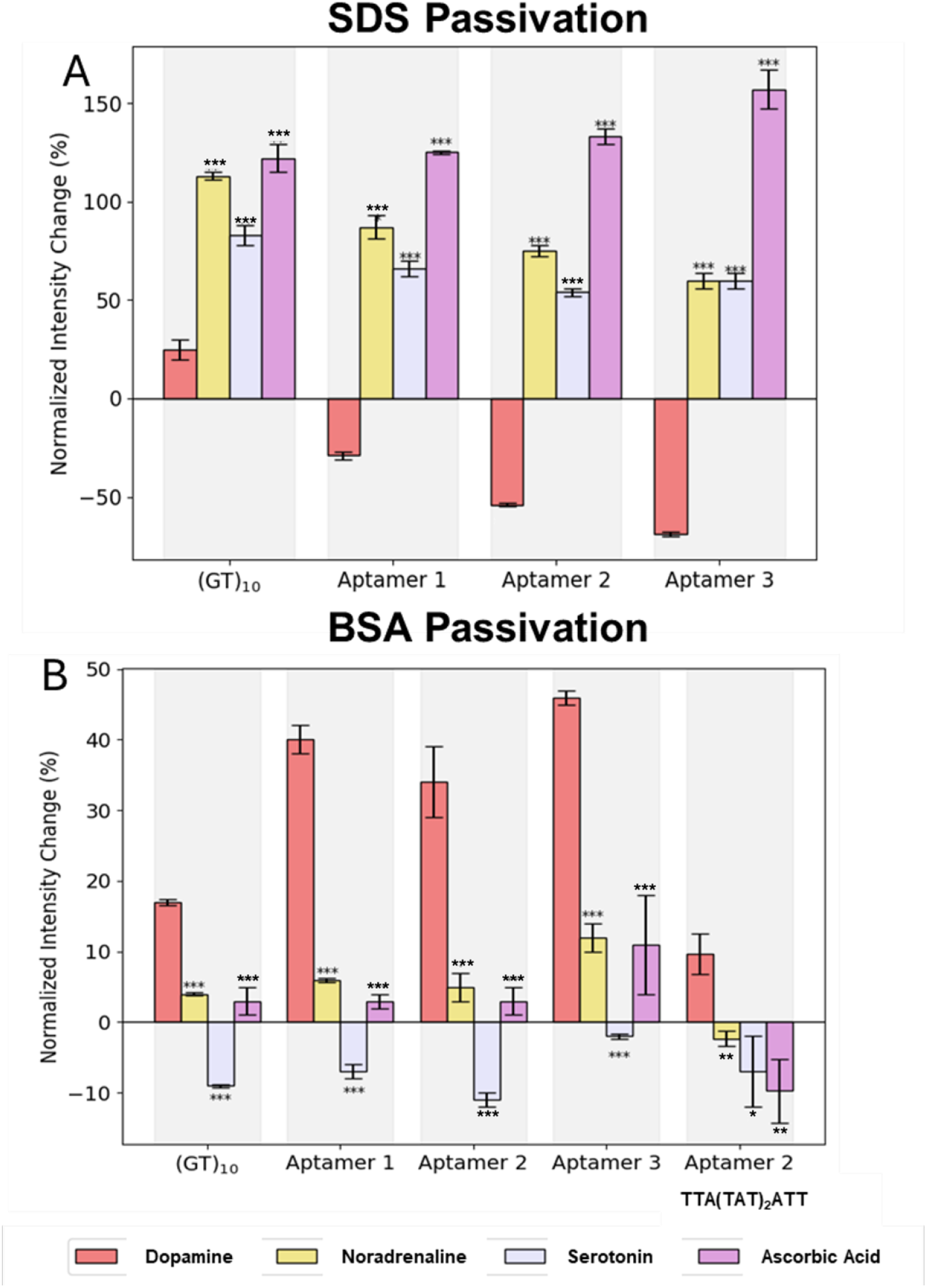
Passivation of sensor constructs improves magnitude of response and selectivity for dopamine. Sensors were passivated with (A) sodium dodecyl sulfate (SDS) and (B) bovine serum albumin (BSA) prior to incubation with 100 nM dopamine or interferents. SWCNT. Significant differences in means were determined for a given DNA - SWCNT hybrid by comparing responses to each interferent with those to dopamine, * p<0.05, ** p<0.01, *** p<0.001.

As we have used BSA in our prior studies with antibody-conjugated SWCNT sensors^45, 46^, we further investigated its potential here to improve the selectivity of ssDNA aptamer-SWCNT sensors. BSA passivated sensors indeed exhibited an improvement in selectivity for dopamine (**Figure 3B**), compared to the results obtained in its absence (**Figure 2B**). The responses of all three aptamer-SWCNT constructs induced a 35-45% increase in brightness in response to dopamine. Both Aptamer 1- and Aptamer 2-SWCNT exhibited negligible (<5%) or slightly negative changes in response to noradrenaline, serotonin, or ascorbic acid. Aptamer 3-SWCNT demonstrated the largest response to dopamine, but also slightly greater responses (∼10%) to noradrenaline and ascorbic acid. Interestingly, the Aptamer 2 conjugate construct exhibited a minimal response (∼10%) to dopamine, but significant decrease in brightness to the other constructs. Thus, this conjugate construct exhibited dampened response but increased selectivity, which is similar to our studies with BSA passivation of antibody-conjugate SWCNT sensors^45, 46^. It is also of note that BSA improved the selectivity of (GT)_10_-SWCNT for dopamine. The response to dopamine did not noticeably change compared to without BSA passivation (**Figure 2A**), however responses to other analytes were negligible or negative. This interesting finding corresponds with prior studies that found the two hydroxy groups on dopamine have some interaction with ssDNA phosphate backbone^24, 47^. Noradrenaline also has two hydroxy groups, though it also has a third hydroxy attached to the alkyl chain, the only difference between it and dopamine, which perhaps interferes with this interaction. Among all the systems evaluated, SWCNT functionalized with aptamers and passivated with BSA exhibited the most favorable combination of selectivity and sensitivity and were therefore selected for subsequent experiments.

We next assessed the response of each BSA-passivated aptamer-SWCNT, and (GT)_10_-SWCNT over time. We found that in each case the intensity of the sensor, in PBS alone and without analytes, decreased slightly over 170 minutes (**Figure 4**). In all cases, the response to dopamine was immediate and substantial, reaching 30-40% increase within 5 minutes for each aptamer-SWCNT and just ∼10% for (GT)_10_-SWCNT. Further, in all cases, the response to noradrenaline and ascorbic acid was a slight decrease in intensity, but less so than PBS alone, and the response to serotonin was a larger decrease than for PBS alone. Nevertheless, it is possible that, since they would be found in the same environment, the response to serotonin and noradrenaline may confound sensor results somewhat.

**Figure 4:**
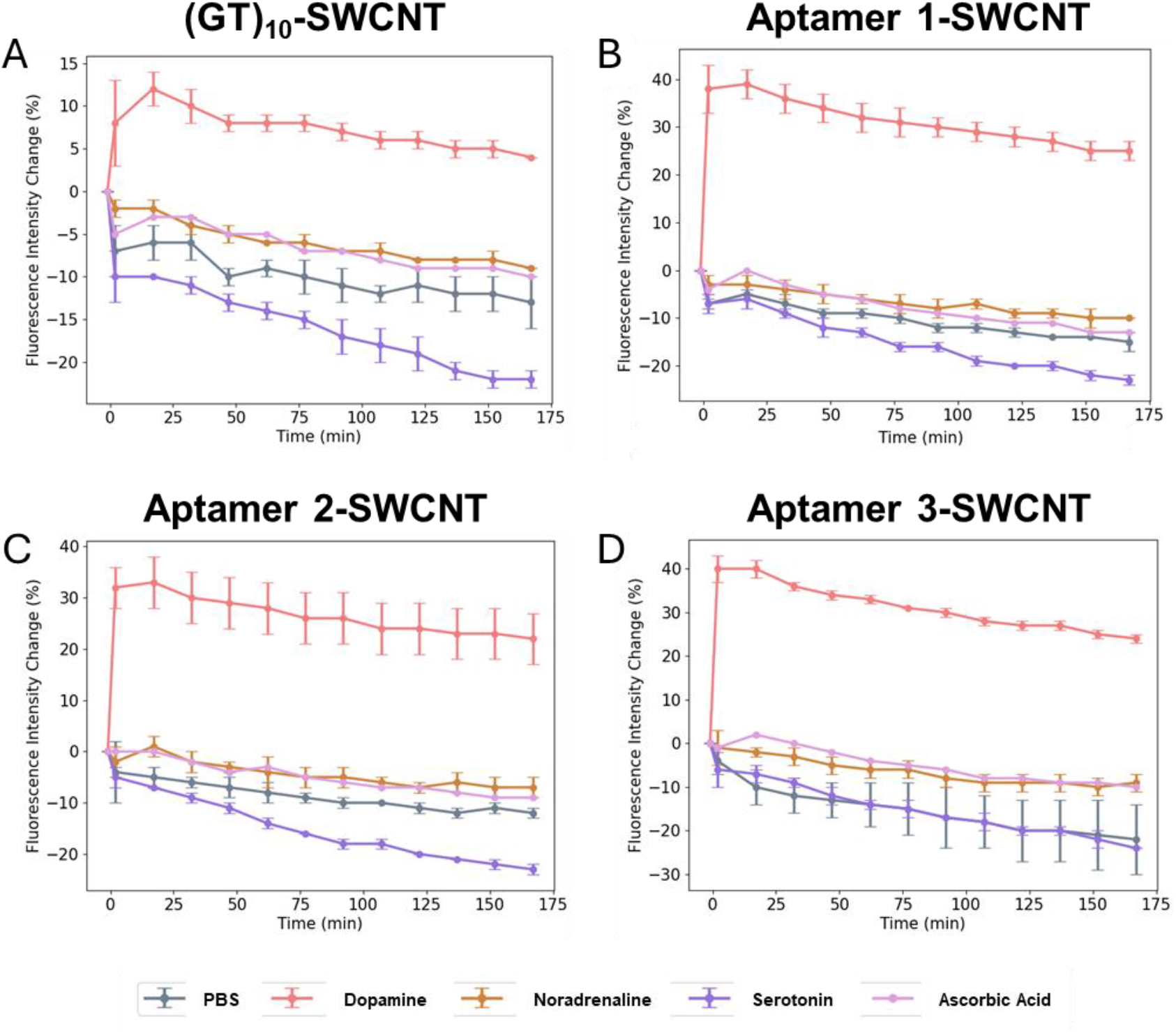
Time-resolved intensity changes in response to analytes for BSA-passivated sensors. Change in intensity in response to 100 nM dopamine and interferents was assessed over 170 minutes.

### Dopamine detection in physiologically-relevant biofluids

Given the improved response of BSA-passivated sensors compared to interferent molecules, we further evaluated their concentration-dependent response. First, a broad concentration range from 1 – 1000 nM of each analyte was tested (**Figure 5A**) to determine the lowest concentration at which the response differed significantly from the baseline, which occurred at 100 nM. We note that the response to 1000 nM (1 µM) dopamine exhibited 75-125% intensity increases for aptamer-SWCNT constructs, while it was approximately 50% increase for (GT)_10_-SWCNT. This is slightly less, but similar to, previous reports of 58-82% increases for (GT)_15_-SWCNT^17^. Based on this threshold, we investigated dose response kinetics to dopamine from 30 nM to 200 nM in PBS (**Figure 5B**). The three aptamers and (GT)_10_ hybrids exhibited a concentration-dependent response, although the response to (GT)_10_ was lower in magnitude for all concentrations tested, while all three aptamers demonstrated approximately similar response at each concentration.

**Figure 5:**
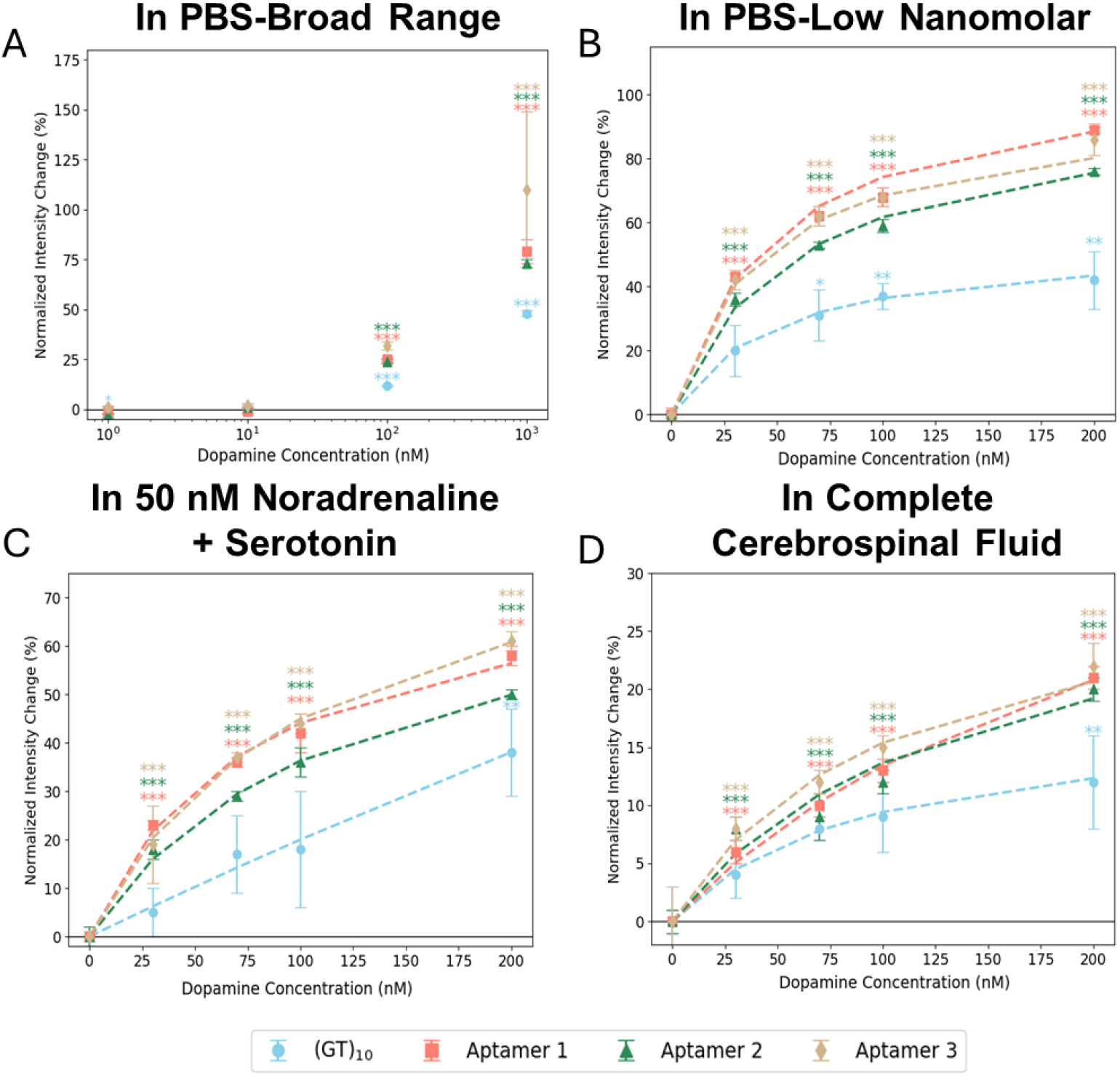
Concentration-dependent response of DNA-SWCNTs to dopamine in buffer and complex media. Normalized intensity changes (%) in the (7,5) SWCNT wrapped with (GT)_10_ and three dopamine aptamers. All were passivated with BSA. Response to (A) a broad range and (B) a narrow range of dopamine concentrations in PBS, (C) dopamine in 50 nM noradrenaline and 50 nM serotonin in PBS, and (D) in aCSF. The dotted line corresponds to the Langmuir fit.

To simulate a more complex environment which may be found *in vitro* or *in vivo*, we investigated sensor response to dopamine in the presence of noradrenaline and serotonin. We performed the same concentration curve as above in a PBS solution containing 50 nM of each interferent. As previously mentioned, dopamine induces an increase in fluorescence intensity, while interferents cause a decrease. In this context, the aim of the experiment was to analyze how the simultaneous presence of serotonin and noradrenaline affects the concentration-dependent response of the SWCNT hybrids. The results (**Figure 5C**) demonstrate that the concentration-dependent behavior was preserved. However, the overall magnitude of response was diminished slightly for all sensor constructs.

To further investigate the response of each sensor construct in artificial *in vivo* conditions, we investigated their concentration-dependent response to dopamine in a simulated cerebrospinal fluid. In these experiments, sensors and dopamine were spiked directly into complete, undiluted aCSF. Again, a concentration-dependent response was observed (**Figure 5D**), however with a substantial decrease in the magnitude of response. It should be noted, however, that the response of each aptamer-SWCNT construct appeared relatively monotonic from 30-200 nM dopamine in aCSF, whereas (GT)_10_-SWCNT responses plateaued at 75 nM. This effect may be attributed to the high ionic strength of the aCSF, which can alter the conformational structure of both (GT)_10_ and the aptamers, thereby modifying their interaction with dopamine^48^

Concentration-response curves were fit to a Langmuir model as in previous reports^49^, treating ssDNA-SWCNT as a single active binding site that reaches saturation (**Table 2**). For all constructs, the dissociation constant (K_d_) was increased in both complex medias, whereas the maximum response was somewhat diminished. The responses to each aptamer-SWCNT construct were generally similar, while that to (GT)_10_-SWCNT had lower K_d_ values, likely attributable to lower maximum intensities (Max NIC). Though, it is clear that some amount of dopamine-ssDNA interaction occurs for (GT)_10_-SWCNT, driven as predicted from prior work by hydroxy interactions with ssDNA phosphates^24, 47^. It is interesting that the model was unable to fit (GT)_10_-SWCNT response in noradrenaline + serotonin, as a linear response was observed. This is likely due to the somewhat less specific nature of interaction of GT-repeats with dopamine. Thus, we find it likely that the increase in response magnitude to dopamine for aptamer-SWCNT constructs was likely partially driven by interact binding site-analyte interactions, as well as nonspecific ssDNA-analyte interactions^24, 47^. It is possible that this response is uniquely transduced by SWCNT due to increased charge transfer from the primary amine on the dopamine alkyl chain.

**Table 2:**
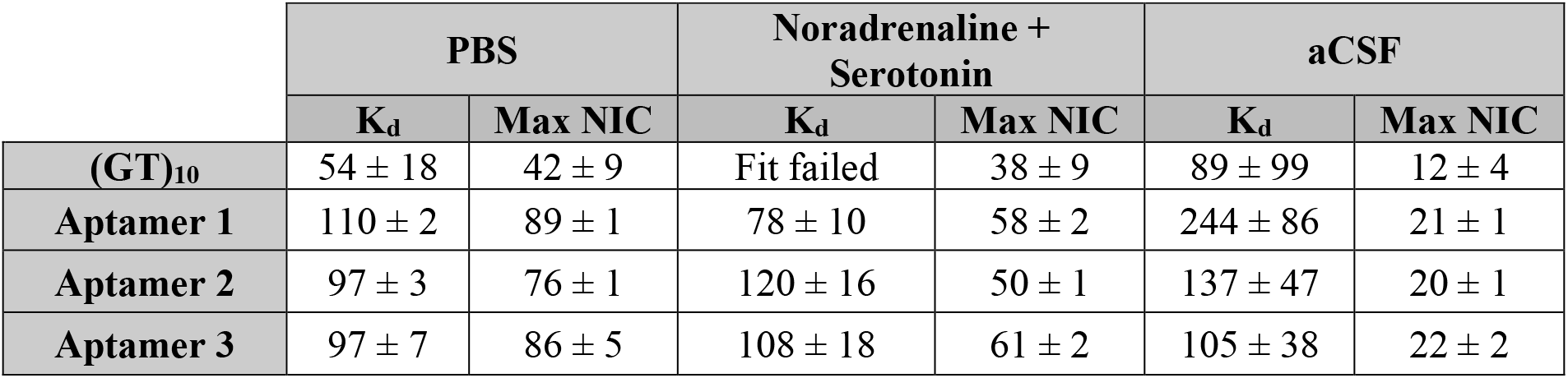
Binding kinetics for each sensor construct and dopamine. Dissociation constant (K_d_) and maximum normalized intensity change (Max NIC) for each followed Langmuir model fit. Concentrations are given in nM.

## Conclusions

In this study, we found that the rational design of SWCNT dopamine sensors with aptamer molecular recognition elements enhances both the magnitude of response and the selectivity compared to current standards. Compared to (GT)_10_–SWCNT sensors, aptamer-SWCNT constructs highlighted the benefits of using chemically evolved recognition elements. However, importantly, this study does indeed demonstrate that there is some selectivity for GT repeats for dopamine especially when passivation agents are used to reduce nonspecific binding, and there may be a general selectivity of ssDNA for dopamine.

We also sought to improve selectivity and robustness of response via tangential sensor design methods. We first investigated thermal + divalent cation treatment of each sensor, which we hypothesized to improve aptamer conformational folding. Indeed, this improved selectivity of all sequences, nonspecific sequences included. We also found that passivation agents enhanced selectivity—specifically SDS induced a “turn-off” sensor, while BSA induced a “turn-on” sensor for each construct. We further investigated BSA as a passivation agent, finding that sensor response, when not normalized, only increased in the presence of dopamine and not other analytes investigated. We found that the magnitude of response for BSA-passivated aptamer-SWCNT sensors was 3-4 times greater than for (GT)_10_. Excitingly, BSA-passivated aptamer-SWCNT constructs maintained a clear concentration-dependent response to dopamine in the presence of 50 nM noradrenaline and serotonin, as well as in simulated cerebrospinal fluid.

Together, these results demonstrate that aptamer-SWCNT sensors may allow for more robust and selective detection of dopamine in both in vitro and in vivo. While using (GT)_10_-SWCNT sensors for dopamine continues to be a valid detection strategy for some studies, the use of GT-rich ssDNA to sense other analytes may hinder its translation. We anticipate that future studies will investigate the use of passivated aptamer-SWCNT sensors beyond the current biological media studies and into dynamic in vitro and in vivo systems.

## Acknowledgements

The authors wish to acknowledge all members of the Williams Lab for discussion and feedback. This work was supported by NIH R35GM142833 and the SUNY Empire Innovation Program (Award #250010) (R. Williams). A. Israel and A. Ryan were supported by a G-RISE Ph.D. traineeship from the National Institutes of Health (T32GM136499).

## References

(1) Speranza, L.; Miniaci, M. C.; Volpicelli, F. The Role of Dopamine in Neurological, Psychiatric, and Metabolic Disorders and Cancer: A Complex Web of Interactions. Biomedicines 2025, 13 (2).

(2) Goldstein, D. S.; Holmes, C.; Lopez, G. J.; Wu, T.; Sharabi, Y. Cerebrospinal fluid biomarkers of central dopamine deficiency predict Parkinson’s disease. Parkinsonism & related disorders 2018, 50, 108–112.

(3) Ceyzériat, K.; Gloria, Y.; Tsartsalis, S.; Fossey, C.; Cailly, T.; Fabis, F.; et al. Alterations in dopamine system and in its connectivity with serotonin in a rat model of Alzheimer’s disease. Brain Communications 2021, 3 (2).

(4) Orhan, F.; Goiny, M.; Becklén, M.; Mathé, L.; Piehl, F.; Schwieler, L.; et al. CSF dopamine is elevated in first-episode psychosis and associates to symptom severity and cognitive performance. Schizophrenia research 2023, 257, 34–40.

(5) Kremer, T.; Taylor, K. I.; Siebourg-Polster, J.; Gerken, T.; Staempfli, A.; Czech, C.; et al. Longitudinal Analysis of Multiple Neurotransmitter Metabolites in Cerebrospinal Fluid in Early Parkinson’s Disease. Movement disorders : official journal of the Movement Disorder Society 2021, 36 (8), 1972–1978.

(6) Choi, Y.; Jeon, C. S.; Kim, K. B.; Kim, H.-J.; Pyun, S. H.; Park, Y. M. Quantitative detection of dopamine in human serum with surface-enhanced Raman scattering (SERS) of constrained vibrational mode. Talanta 2023, 260.

(7) Colniţă, A.; Marconi, D.; Toma, V. A.; Brezeştean, I.-A.; Suciu, M.; Ciorîţă, A.; et al. SERS detection of dopamine in artificial cerebrospinal fluid and in Parkinson’s disease-induced mouse cortex using a hybrid ZnO@Ag nanostructured platform. Microchemical Journal 2024, 206.

(8) Ma, X.; Wu, Y.; She, J.; Zhao, A.; Yang, S.; Yang, X.; et al. On-chip electrochemical sensing of neurotransmitter in nerve cells by functionalized graphene fiber microelectrode. Sensors and Actuators B: Chemical 2022, 365.

(9) Zhang, C.; Chen, T.; Ying, Y.; Wu, J. Detection of Dopamine Based on Aptamer-Modified Graphene Microelectrode. Sensors 2024, 24 (9).

(10) Magar, H. S.; Duraia, E.-s. M.; Hassan, R. Y. A. Dopamine fast determination in pharmaceutical products using disposable printed electrodes modified with bimetal oxides carbon nanotubes nanocomposite. Scientific Reports 2025, 15.

(11) O’Connell, M. J.; Bachilo, S. M.; Huffman, C. B.; Moore, V. C.; Strano, M. S.; Haroz, E. H.; et al. Band gap fluorescence from individual single-walled carbon nanotubes. Science 2002, 297.

(12) Cohen, Z.; Williams, R. M. Single-Walled Carbon Nanotubes as Optical Transducers for Nanobiosensors In Vivo. ACS Nano 2024, 18 (52).

(13) Ryan, A. K.; Rahman, S.; Williams, R. M. Optical Aptamer-Based Cytokine Nanosensor Detects Macrophage Activation by Bacterial Toxins. ACS Sensors 2024, 9 (7).

(14) Zanetti, J. K.; Stefoni, M. C.; Ferraz, C.; Ryan, A.; Israel, A.; Williams, R. M. A near-infrared fluorescent aptananosensor enables selective detection of the stress hormone cortisol in artificial cerebrospinal fluid. Sensors & Diagnostics 2025, 4 (12), 1103–1113.

(15) Dinarvand, M.; Neubert, E.; Meyer, D.; Selvaggio, G.; Mann, F. A.; Erpenbeck, L.; et al. Near-Infrared Imaging of Serotonin Release from Cells with Fluorescent Nanosensors. Nano Letters 2019, 19 (9), 6604–6611.

(16) Basu, S.; Hendler-Neumark, A.; Bisker, G. Role of Oxygen Defects in Eliciting a Divergent Fluorescence Response of Single-Walled Carbon Nanotubes to Dopamine and Serotonin. ACS Nano 2024, 18 (50).

(17) Kruss, S.; Landry, M. P.; Ende, E. V.; Lima, B. M. A.; Reuel, N. F.; Zhang, J.; et al. Neurotransmitter Detection Using Corona Phase Molecular Recognition on Fluorescent Single-Walled Carbon Nanotube Sensors. Journal of American Chemical Society 2013, 136 (2).

(18) Basu, S.; Bisker, G. Near-Infrared Fluorescent Single-Walled Carbon Nanotubes for Biosensing. Small 2025, 21 (26).

(19) Nißler, R.; Kurth, L.; Li, H.; Spreinat, A.; Kuhlemann, I.; Flavel, B. S.; et al. Sensing with Chirality-Pure Near-Infrared Fluorescent Carbon Nanotubes. Analytical Chemistry 2021, 93 (16).

(20) Liu, X.; Chen, J.; Wang, H.; Lambert, B.; Boghossian, A. A. Cation Pretreatment Enables the Saline Stability of a Near-Infrared Sensor for Dopamine. ACS Bio & Med Chem Au. 2025, 5 (1), 166–174.

(21) Beyene, A. G.; Alizadehmojarad, A. A.; Dorlhiac, G.; Goh, N.; Streets, A. M.; Král, P.; et al. Ultralarge Modulation of Fluorescence by Neuromodulators in Carbon Nanotubes Functionalized with Self-Assembled Oligonucleotide Rings. Nano Letters 2018, 18 (11), 6995–7003.

(22) Hill, B. F.; Mohr, J. M.; Sandvoss, I. K.; Gretz, J.; Galonska, P.; Schnitzler, L.; et al. Ratiometric near infrared fluorescence imaging of dopamine with 1D and 2D nanomaterials. Nanoscale 2024, (39).

(23) Dinarvand, M.; Elizarova, S.; Daniel, D. J.; Kruss, D. S. Imaging of Monoamine Neurotransmitters with Fluorescent Nanoscale Sensors. ChemPlusChem 2020, 1465–1480.

(24) Mann, F. A.; Herrmann, N.; Meyer, D.; Kruss, S. Tuning Selectivity of Fluorescent Carbon Nanotube-Based Neurotransmitter Sensors. Sensors 2017, 17 (7).

(25) Jin, H.; Heller, D. A.; Kalbacova, M.; Kim, J.-H.; Zhang, J.; Boghossian, A. A.; et al. Detection of single-molecule H2O2 signalling from epidermal growth factor receptor using fluorescent single-walled carbon nanotubes. Nature nanotechnology 2010, 5 (4), 302–309.

(26) Cohen, Z.; Alpert, D. J.; Weisel, A. C.; Ryan, A.; Roach, A.; Rahman, S.; et al. Noninvasive Injectable Optical Nanosensor-Hydrogel Hybrids Detect Doxorubicin in Living Mice. Advanced Optical Materials 2024, 2303324.

(27) Harvey, J. D.; Williams, R. M.; Tully, K. M.; Baker, H. A.; Shamay, Y.; Heller, D. A. An in vivo nanosensor measures compartmental doxorubicin exposure. Nano letters 2019, 19 (7), 4343–4354.

(28) Ranjbar-Slamloo, Y.; Fazlali, Z. Dopamine and Noradrenaline in the Brain, Overlapping or Dissociate Functions? Front. Mol. Neurosci. 2020, 12.

(29) Liu, X. L. a. J. Biosensors and sensors for dopamine detection. VIEW 2020, 2.

(30) Miled, S. D. N. a. P. K. a. P. X. a. J. M. a. P. D. K. a. E. B. a. M. B. a. A. A Review of Neurotransmitters Sensing Methods forNeuro-Engineering Research. Applied Sciences 2019, 9.

(31) Ao, G.; Streit, J. K.; Fagan, J. A.; Zheng, M. Differentiating left-and right-handed carbon nanotubes by DNA. Journal of the American Chemical Society 2016, 138 (51), 16677–16685.

(32) Zheng, M.; Jagota, A.; Semke, E. D.; Diner, B. A.; McLean, R. S.; Lustig, S. R.; et al. DNA-assisted dispersion and separation of carbon nanotubes. Nature materials 2003, 2 (5), 338–342.

(33) ZhaoJing-Wen, T.; Zhang, W.-S.; Chen, X. Z.-P.; Tang, H.; Jiang, J.-H. Development of Dual-Nanopore Biosensors for Detection of Intracellular Dopamine and Dopamine Efflux from Single PC12 Cell. Analytical Chemistry 2022, 94 (45).

(34) Wu, G.; Zhang, N.; Matarasso, A.; Heck, I.; Li, H.; Lu, W.; et al. Implantable Aptamer-Graphene Microtransistors for Real-Time Monitoring of Neurochemical Release in Vivo. Nano Letters 2022, 22 (9).

(35) Xu, Y.; Hun, X.; Liu, F.; Wen, X.; Luo, X. Aptamer biosensor for dopamine based on a gold electrode modified with carbon nanoparticles and thionine labeled gold nanoparticles as probe. Microchimica Acta 2015, 182.

(36) Hebling, C. M.; Thompson, L. E.; Eckenroad, K. W.; Manley, G. A.; Fry, R. A.; Mueller, K. T.; et al. Sodium Cholate Aggregation and Chiral Recognition of the Probe Molecule (R,S) 1,1′-binaphthyl-2,2′ diylhydrogenphosphate (BNDHP) Observed by 1H and 31P NMR Spectroscopy. Langmuir 2009, 24 (24), 13866–13874.

(37) Andy Coffey, A. T., Inc. Size Exclusion Chromatography in the Presence of an Anionic Surfactant; 2017.

(38) Gaikwad, P.; Rahman, N.; Parikh, R.; Crespo, J.; Cohen, Z.; Williams, R. M. Optical Nanosensor Passivation Enables Highly Sensitive Detection of the Inflammatory Cytokine Interleukin-6. ACS Appl Mater Interfaces 2024, 16 (21), 27102–27113.

(39) Channer, B.; Matt, S. M.; Nickoloff-Bybel, E. A.; Pappa, V.; Agarwal, Y.; Wickman, J.; et al. Dopamine, Immunity, and Disease. Pharmacological Reviews 2023, 75 (1), 62–158.

(40) Williams, G. V.; Millar, J. Concentration-dependent actions of stimulated dopamine release on neuronal activity in rat striatum. Neuroscience 1990, 39 (1), 1–16.

(41) Nakatsuka, N.; Abendroth, J. M.; Yang, K.-A.; Andrews, A. M. Divalent Cation Dependence Enhances Dopamine Aptamer Biosensing. 2021, 13 (8), 9425–9435.

(42) Rebecca L. Pinals, F. L., Darwin Yang, Nicole Navarro, Sanghwa Jeong, John E. Pak, Lili Kuo, Yung-Chun Chuang, Yu-Wei Cheng, Hung-Yu Sun, Markita P. Landry. Rapid SARS-CoV-2 Spike Protein Detection by Carbon Nanotube-Based Near-Infrared Nanosensors. Nano Letters 2021, 21 (5), 2272–2280.

(43) Darwin Yang, S. J. Y., Jackson Travis Del Bonis-O’Donnell, Rebecca L. Pinals, Markita P. Landry. Mitigation of Carbon Nanotube Neurosensor Induced Transcriptomic and Morphological Changes in Mouse Microglia with Surface Passivation. ACS Nano 2020, 14 (10), 13794–13805.

(44) Y.L. Jeyachandran, J. A. M., E. Mielczarski, B. Rai. Efficiency of blocking of non-specific interaction of different proteins by BSA adsorbed on hydrophobic and hydrophilic surfaces. Journal of Colloid and Interface Science 2010, 341 (1), 136–142.

(45) Williams, R. M.; Lee, C.; Galassi, T. V.; Harvey, J. D.; Leicher, R.; Sirenko, M.; et al. Noninvasive ovarian cancer biomarker detection via an optical nanosensor implant. Science Advances 2018, 4 (4).

(46) Williams, R. M.; Lee, C.; Heller, D. A. A Fluorescent Carbon Nanotube Sensor Detects the Metastatic Prostate Cancer Biomarker uPA. ACS Sensors 2018, 3 (9), 1838–1845.

(47) Kruss, S.; Salem, D. P.; Vuković, L.; Lima, B.; Ende, E. V.; Boyden, E. S.; et al. High-resolution imaging of cellular dopamine efflux using a fluorescent nanosensor array. PNAS 2017, 114 (8), 1789–1794.

(48) Luo, Y.; Wu, N.; Niu, L.; Hao, P.; Sun, X.; Chen, F.; et al. Ionic Strength-Mediated “DNA Corona Defects” for Efficient Arrangement of Single-Walled Carbon Nanotubes. Advanced Science 2024, 11 (15).

(49) Santos, T.; Lopes-Nunes, J.; Alexandre, D.; Miranda, A.; Figueiredo, J.; Silva, M. S.; et al. Stabilization of a DNA aptamer by ligand binding. Biochimie 2022, 200, 8–18.

