## Supplemental Figures for "Selective and robust dopamine detection is enabled by aptamer-SWCNT optical sensors in physiological media"

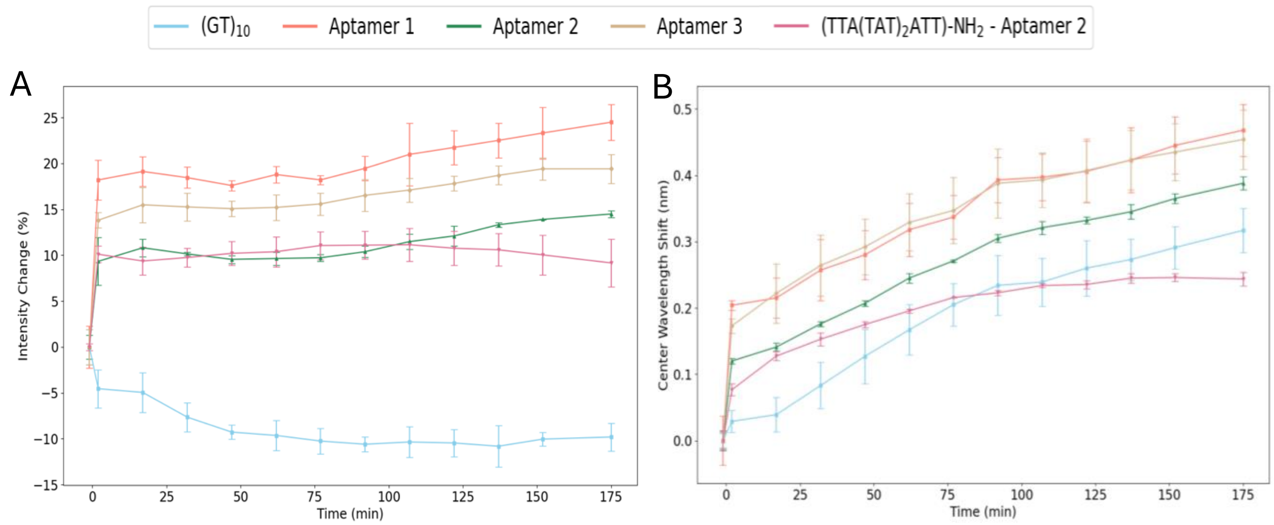


**Supplementary Figure S1: Fluorescence changes relative to untreated controls over time in response to dopamine.** (A) Intensity changes (%) and (B) center wavelength shift, in the (7,5) peak of SWCNTs wrapped with $(GT)_{10}$, three dopamine aptamers, and the $(TTA(TAT)_2ATT)\text{-}NH_2$ aptamer 2 conjugate. The response was studied with dopamine at a final concentration of 100 nM.

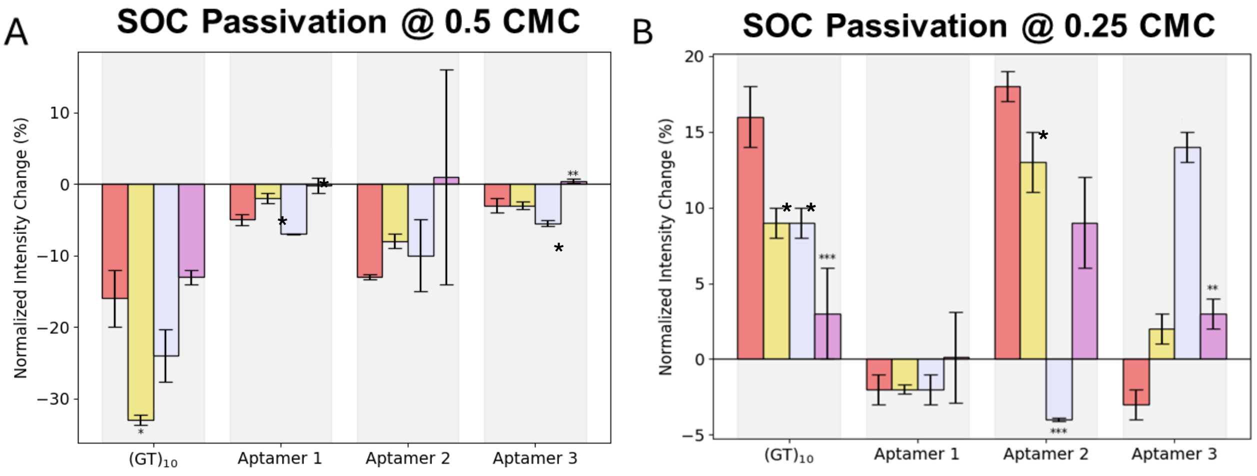


**Supplementary Figure S2: Sensor response to dopamine and interferents in the presence of SOC as a passivation agent.** SWCNT wrapped with $(GT)_{10}$, dopamine aptamers, or Aptamer 2-conjugate were passivated with (A) Sodium Cholate (SOC) at 0.5 critical micelle concentration (CMC) and (B) 0.25 CMC. Significance of response compared to dopamine was determined * $p<0.05$, ** $p<0.01$, *** $p<0.001$.

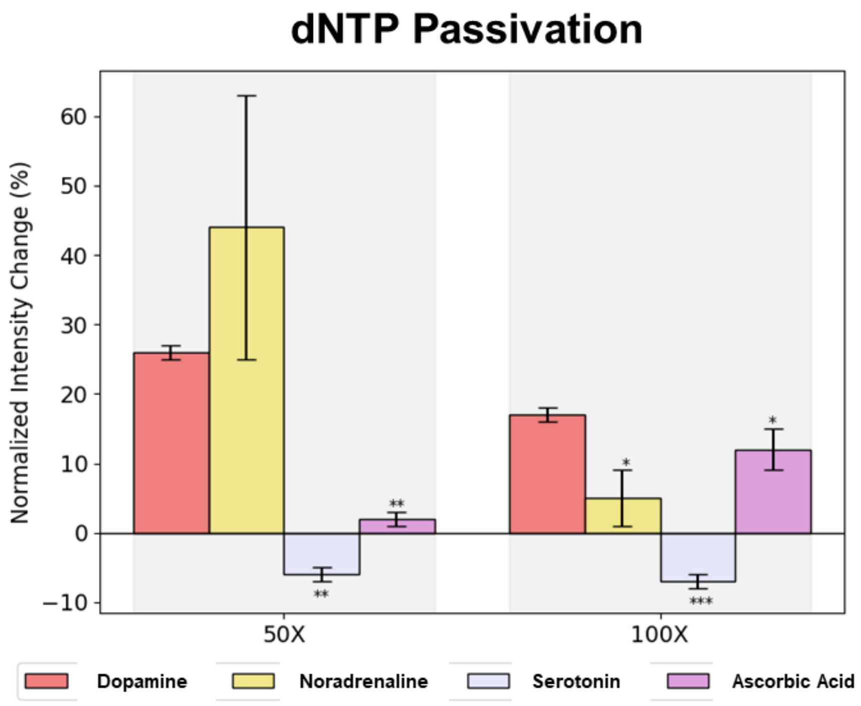


**Supplementary Figure S3: Sensor response to dopamine and interferents in the presence of dNTPS as a passivation agent.** SWCNT wrapped with dopamine aptamer 2. dNTPs at a 50x and 100x mass ratio to SWCNT were added prior to incubation with analytes at 100 nM. Significance of response compared to dopamine was determined * $p<0.05$, ** $p<0.01$, *** $p<0.001$.
